# Humanized mice for sustained *Plasmodium vivax* blood-stage infection and transmission

**DOI:** 10.1101/2022.03.22.485265

**Authors:** Camilla Luiza-Batista, Sabine Thiberge, Malika Serra-Hassoun, Flore Nardella, Aurélie Claes, Vanessa C. Nicolete, Pierre-Henri Commère, Liliana Silva-Mancio, Marcelo U. Ferreira, Artur Scherf, Sylvie Garcia

## Abstract

*Plasmodium vivax* is the most widespread human malaria parasite ^1^. The presence of extravascular reservoirs ^2^, the early circulation of infective stages (gametocytes), and relapsing infections arising from dormant liver stages ^3^ render this parasite particularly difficult to control and eliminate ^4^. Experimental research is limited by the lack of a continuous culture in vitro system that fulfills the parasite’s needs,^5^ namely its tropism for immature CD71^+^ red blood cells (RBCs). ^5^ Here, we report a humanized mice model, which upon engraftment of human hematopoietic progenitor and stem cells (HPSCs), exhibits efficient human erythropoiesis. Humanized HIS-HEry mice inoculated with cryopreserved *P. vivax* samples sustain long-lasting asexual parasite multiplication within CD71^+^ human RBCs and differentiation into mature gametocytes that can be efficiently transmitted to *Anopheles* mosquitoes, leading to formation salivary-gland sporozoites. Blood stages can be sequentially transferred to uninfected humanized mice by injection of fresh or frozen infected bone marrow cells, providing a unique murine model for the long-term maintenance of *P. vivax* isolates. This work offers a novel experimental platform to investigate the biology of RBC invasion and intraerythrocytic *P. vivax* development in vivo and evaluate new interventions against this elusive human parasite.

## Main

Malaria poses a major public health burden worldwide, with 229 million clinical cases and 384,000 deaths in 2019 according to the WHO **^1^**. Two *Plasmodium* species are responsible for the vast majority of human infections: *P. falciparum,* which predominates in Africa and causes more-severe disease, and *P. vivax*, the globally widespread parasite associated with debilitating illness and occasional severe complications **^4^**. Malaria elimination is still hampered by drug resistance and the lack of an efficient vaccine. The dormant liver stages in *P. vivax* (so called hypnozoites) **^3^** together with extravascular parasite reservoirs such as bone marrow (BM) **^6^** represent a major challenge for eradication strategies. While the design of new control strategies requires a good understanding of *P. vivax* biology and host-parasite interactions, a major roadblock is the lack of an in vitro continuous culture that sustains the parasite erythrocytic cycle **^7^.** This is in part due to the parasite’s tropism for CD71^+^ immature RBCs that are rare in the peripheral blood (PB) **^5^**. Humanized mice able to maintain *P. vivax* infection through *de novo* production of CD71^+^ human RBCs might help to circumvent this drawback, but current models are limited by poor human erythropoiesis and do not sustain *P. vivax* blood-stage infection.

### HIS-HEry mice support human erythropoiesis

CH1-2hSa mice are alymphoid RAG-γc- that express a human SIRP receptor, the receptor for the “don’t eat me” signal regulatory protein CD47, under the *CSF1R*-promoter, and HLA-A2 and HLA-DR1 molecules in place of murine MHC **^8^.** Upon neonatal intra-hepatic engraftment with human cord blood (CB) CD34^+^ cells, CH1-2hSa mice presented robust levels of human leucocytes with approximately 30% of human CD45+ cells in the PB 4-5 months after engraftment **^8^**. In addition, we showed that CH1-2hSa mice were able to mount human functional HLA-restricted T cell immune response. However, virtually no human erythrocytes (CD235a^+^) were present in the bone marrow (BM) of the chimeras (Figure 1a, left panel), consistent with the impaired erythropoiesis previously described in other humanized mice models **^9^**. To improve human red blood cell (RBC) production in CH1-2hSa chimeras, we introduced the hypomorphic KITW41 mutation of the murine Stem Cell Factor Receptor (SCF-R or cKit). By decreasing murine hematopoietic cell fitness and providing available hematopoietic niches for human engrafted CD34^+^ cells **^9–11^**, cKITW41 expression dramatically increased the percentage of CD235a^+^ human red cells in the BM of CH1-2hSa chimeras (Figure 1a, right panel), resulting in robust human erythropoiesis (about 25% of the RBCs being of human origin) (Fig. 1c). As expected for developing erythrocytes, 70% of the human bone marrow (BM) CD235a^+^ erythroid stages expressed CD71 (Fig. 1b, c), a key receptor for *P. vivax* RBC invasion **^5^**. The co-expression profile of these two markers and proportion of CD71^+^ CD235a^+^ cells were consistent with what has been observed in human BM (Fig. 1b, c) ^12^. The physiological maturation of human RBCs in the murine environment was further confirmed by the relative expression of CD49d and Band3 (Extended data Fig. 1a), induction of human adult hemoglobin expression (Extended data Fig. 1c) and the expression levels of different markers of erythropoiesis – namely, CD235a, CD36, CD44, CD234 (DARC) (another entry receptor for *P. vivax*) ^13^, and CD71 – by human CD235a^+^ erythroid cells in the chimeric BM (Extended data Fig. 1c). We named this new humanized mouse model HIS-HEry for <u>H</u>uman Immune <u>S</u>ystem <u>H</u>uman <u>Ery</u>throcytes. In contrast to the BM compartment, human RBCs were detected only at low levels in the peripheral blood (PB), likely due to destruction by peripheral murine macrophages as reported elsewhere ^9,14^ (Extended data Fig. 1d). The injection of clodronate liposomes, which deplete mouse macrophages, significantly raised the presence of human RBC in the peripheral blood to approximatively 3% of total circulating RBC (Extended data Fig. 1d).

**Fig. 1.**
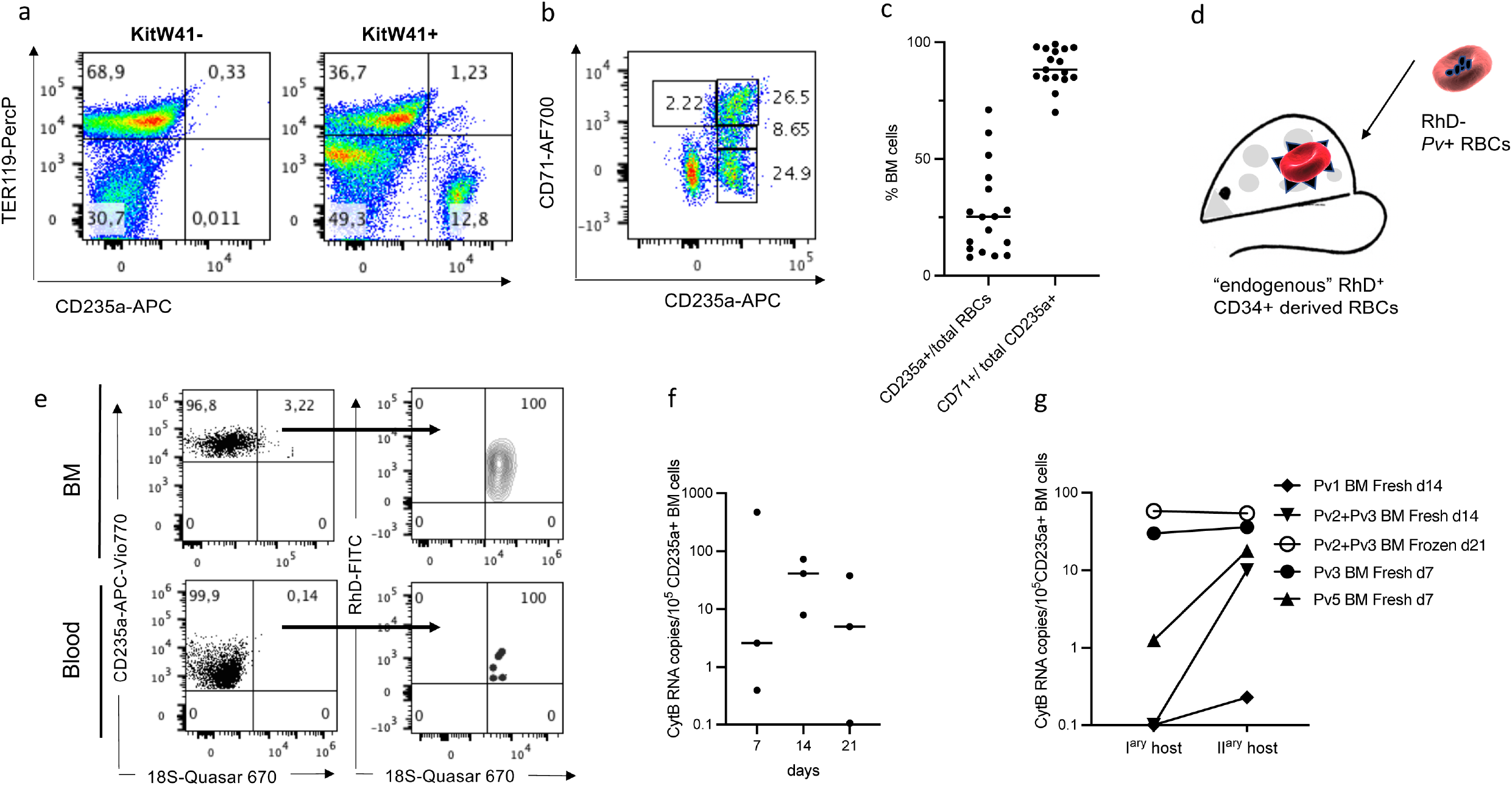
HIS-HEry chimeras support human erythropoiesis and *de novo* infection by *P. vivax*. Twenty to 24 weeks old HIS-HEry (KitW41+) BM cells were analyzed by flowcytometry for: **a-** Respective representation of human CD235a+ versus murine TER119 RBCs in KitW41^−^ and KITW41^+^ chimeras. **b-** Representative co-expression of CD71 relative to CD235a marker by TER119^−^ Ly5.2^−^ (murine leucocytes) CD45^−^ (human leucocytes) cells. **c-** Distribution of human CD235^+^ RBCs among total RBCs (left) and the percentage of total CD71^+^ cells among total human CD235^+^ RBCs (right) in HIS-HEry BM chimeras (*n*=17). **d-e-** HIS-HEry mice reconstituted with RhD^+^ CD34^+^ cord blood cells were infected with *P. vivax* (isolate Pv1)-infected RBCs from RhD^−^ patient and human RBCs processed by FlowFISH at day7 post infection. **e-** *P. vivax* specific 18S^+^ parasitized cells were gated among CD235a^+^ BM (upper left dot plot) and PB (lower left dot plot) human cells, and parasitized 18S^+^ cells was analyzed among human RhD^+^ versus RhD^−^ CD235a^+^ BM (upper right dot plot) and PB (lower right dot plot) RBCs. **f-** HIS-HEry chimeras *n*=8 mice reconstituted with the CD34^+^ cells from the same CB donor were infected with a mixture of *P. vivax* isolates Pv2+Pv3 (see table). Numbers of *P. vivax* CytB RNA copies per 10^5^ CD235a^+^ BM cells monitored by qRT-PCR at day7, 14 and 21 post-infection are plotted. Each dot represents one individual mouse. Mean CytB RNA copy numbers are: day 7= 2.59, day 14= 41.31 and day 21= 5.01. This experiment is representative of 3 independent experiments. **g-** BM human RBCs from infected HIS-HEry mice were adoptively transferred into secondary naïve HIS-HEry recipient mice. Numbers of *Plasmodium* CytB RNA copies per 10^5^ human BM RBCs cells were compared between donor cells and secondary host BM 7 days after transfer. Each type of dots represents individual BM inoculum (left) and the corresponding BM from secondary mice (right). The type of BM donor cells (*Pv* isolate, day post-infection, fresh or frozen) is indicated on the figure. *n* = 5 mice.

### HIS-HEry mice support intraerythrocytic *P. vivax* growth

We next investigated whether CD71+ human reticulocytes from HIS-HEry mice could support *P. vivax* infection and multiplication. We used cryopreserved *P. vivax*-infected blood derived from patients from Brazil, with parasitemia levels ranging from 0.015% to 0.045% (SupplementalTable). Infected Rhesus D allele-negative (RhD^−^) RBCs from patient Pv1 were thawed and injected intravenously into HIS-HEry mice reconstituted with cord blood HPSCs from a RhD+ donor (Fig. 1d). Mice were maintained under clodronate liposomes throughout the experiment. To distinguish between RhD+ host HIS-HEry mice RBCs (“endogenous “) and RhD-infected RBCs from the original parasitized human donor, we use a new Flowcytometric RNA FISH method we recently described **^15^** that allowed us to detect both RhD surface expression and infection status of RBCs through *P. vivax* 18S rRNA expression. Remarkably, at day 7 post infection, we observed infected RhD+ reticulocytes produced by HER-HEry mice in both the BM and the peripheral blood, consistent with RBC invasion events in the murine host (Fig. 1e and Extended data Fig.2a), but virtually no infected RhD-RBCs from the parasitized human donor (phenotype: RhD-18S+ CD235a+ RBCs) (Fig. 1e).

We then assessed whether *P. vivax* infection in HIS-HEry mice can be maintained for extended periods of time. We measured *P. vivax*-specific transcript levels over time in the BM of infected mice reconstituted with the same CB cells and infected with the same a mixture of Pv2+Pv3 isolates. *P. vivax* RNA levels were variable within the experimental group of mice, but remained detectable over 21 days post-infection, with a raise 2 weeks after infection (Fig. 1f).

Importantly, the adoptive transfer of fresh and frozen BM cells from previously infected mice into naïve secondary HIS-HEry mice allowed to establish an infection (Fig. 1g). We used BM samples collected at various time points after injection of HIS-HEry mice with different isolates. Four out of five secondary transfers resulted in *P. vivax* parasitemia levels 7 days after inoculation that were higher (based on the CytB transcript levels) than in the original BM cells inoculated (Fig. 1g). The most striking increase was observed for two BM samples with very low parasitemia prior to injection in secondary host mice (Fig. 1g). In addition, frozen BM transfer resulted in the maintenance of parasite load in the secondary host mouse, demonstrating that freezing *P. vivax*-infected BM cells provides a method to store parasite lines that infect HIS-HEry mice. Our results demonstrate that long-term *in vivo* propagation of *P. vivax* blood stages is now possible in a murine model. Moreover, successive passages of *P. vivax* in HIS-HEry mice may eventually select for parasites with increased multiplication rate.

### Mature gametocytes in HIS-HEry mice

To test whether sexual differentiation of *P. vivax* occurs in the HIS-HEry chimeras, we determined by FlowFISH analysis the distribution of gametocytes versus asexual parasites in both BM and PB of HIS-HEry mice using a set of RNA probes specific for the mature gametocyte marker Pvs25 ^16^ (Fig. 2a). We observed a larger proportion of gametocytes in the BM (64% ± 17) versus PB (18% ± 9) 7 days after infection (Fig. 2b). Given that no infected RBCs originating from the injected isolate was detected at day 7 post-infection (Fig. 1e), we conclude that sexual commitment occurs in human erythrocytes produced *de novo* in the chimeric mice. This is also supported by the FlowFISH detection of Pvs25^+^ 18S^+^ gametocytes within RhD^+^ CD235a^+^ BM RBCs (Extended data Fig. 2a). Of note, the observation that BM nucleated RBCs are infected by gametocytes indicates that erythroblasts not only support asexual parasite proliferation but also sexual differentiation (Extended data Fig. 2a). In addition, we found that Pvs25 transcript levels tended to increase between day 1 and day 7 after infection in the BM of HIS-HEry mice, although this difference was not statistically significant (Extended data Fig. 2b). These data are in favor of the preferential residence of gametocytes within the BM in our model, similar to finding reported from human studies **^16,17^**.

**Fig. 2.**
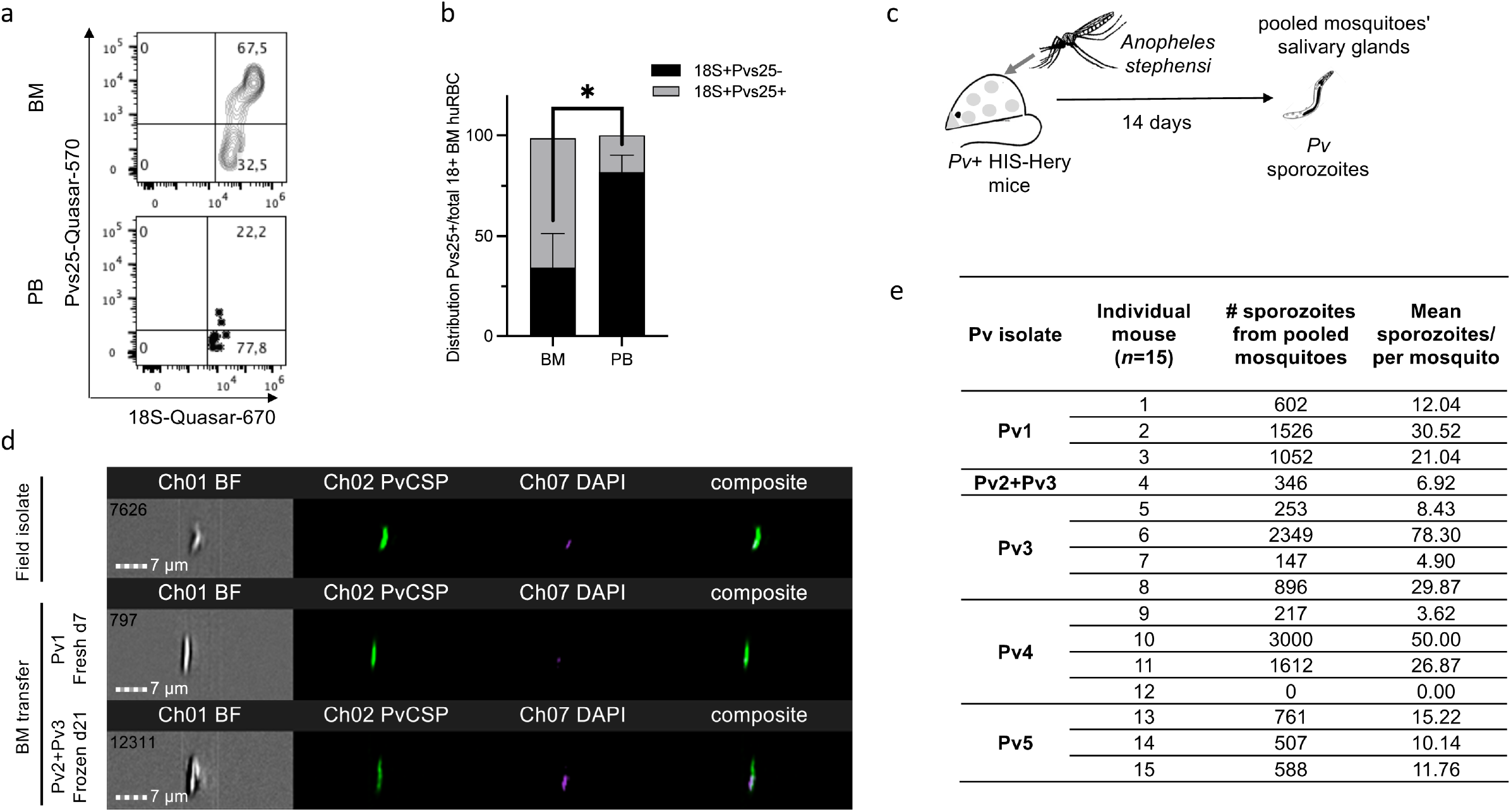
HIS-HEry chimeras support sexual differentiation, *P. vivax* transmission to *Anopheles* and sporozoites production. Representative FlowFISH **a-** plots of *P. vivax* gametocyte specific Pvs25 RNA expression within total *P. vivax*-18S+ parasitized human CD235a^+^ cells in the BM (upper panel) and PB (lower panel) of HIS-HEry 7 days after infection with the Pv1 isolate. **b-** distribution of Pvs25+ 18S+ gametocytes versus Pvs25^−^ 18S^+^ asexual blood stages among BM human CD235a^+^ RBCs. *n* = 5 mice. Data were analyzed with paired t-test P value= 0.0224 (BM Pvs25+ mean= 64,225 ± 17.46; PB Pvs25+ mean= 18.325 ± 8,51). **c-** HIS-HEry infected chimeras were bitten by *Anopheles stephensi* mosquitoes and the transmission efficiency was assessed 14 days later by sporozoite production in the invertebrate host salivary glands. **d**-Representative ImageStream profiles of salivary gland sporozoites stained by FlowFISH with *P. vivax* circumsporozoite protein (PvCSP) in green (Chanel 2 for FITC), nuclei colored with DAPI (Chanel 7) and composite image with colocalization of nuclei and surface PvCSP (right merge). The different images were obtained from: frozen patient field blood isolate (upper), mosquitoes fed on day7 Pv1-infected HIS-HEry mice (middle) and mosquitoes fed on day7 HIS-HEry mice infected with frozen BM from day 21-infected mice (see Fig.2b) (bottom). **e-** Numbers of sporozoites from salivary glands from pooled mosquitoes and estimated numbers of sporozoites per mosquito are shown for each individual mouse (*n* = 15) from each infection experiment with defined *P. vivax* isolates (*n* = 5).

### *P. vivax* sporozoite production

Given the presence of gametocyte stages in the PB of the *P. vivax*-infected HIS-HEry mice, we aimed to assess their viability and maturation by performing transmission assays. HIS-HEry mice infected with different *P. vivax* isolates were used to feed *Anopheles stephensi*. Fourteen days after exposure to mice, salivary glands from mosquitoes were dissected and the presence of sporozoites was analyzed using specific antibodies against the major surface circumsporozoite protein (CSP) (Fig. 2c, 2d). Mosquito infection was highly efficient as gametocytes from 14 of 15 infected mice were transmitted to mosquitoes via blood feeding and originated sporozoites in the mosquito host (Fig. 2e). The numbers of sporozoites derived from infection of individual mice (i.e. from 30-60 mosquitoes per mouse) varied widely according to the isolates, mouse and number of blood-fed mosquitoes, ranging from 147 to 3000 sporozoites. Importantly, the transmission of gametocytes to mosquitoes from secondary infected mice engrafted with frozen BM cells 21 days after infection of donor mice was also successful (Fig. 2e, (Pv2+Pv3), 346 sporozoites), validating the model to perform long-term transmission experiments with the same clinical isolates.

## Discussion

The generation of a novel humanized mouse model able to produce high levels of human erythropoiesis overcomes a major obstacle in malaria research. We explored the potential of HIS-HEry humanized mice for *in vivo* long-term proliferation of *P. vivax*. We obtained *P. vivax* blood stage growth and proliferation for at least three weeks after infection and established a robust protocol to transfer fresh or frozen infected BM cells from HIS-HEry to maintain *P. vivax* infection in a secondary uninfected mouse.

Our HIS-HEry model enables the use of clinical isolates of *P. vivax* in long-term in vivo experiments and mimics several of the important features observed in *P. vivax*-infected humans. First, both sexual and asexual *P. vivax* blood stages were found in the BM, as it was described in BM from patient’s biopsies **^17^**. Second, the high levels of sexually committed parasites found in BM, as compared to the PB, is consistent with previous results obtained from BM biopsies from *P. vivax* infected patients **^16,17^** and supports the hypothesis that BM may represent a niche and a reservoir for sexual stage parasites. Third, we demonstrated the generation of *de novo* gametocytes a few days after mouse infection, confirming that sexual differentiation is faster during *P. vivax* infection than *P. falciparum* infection **^3,18^**. Fourth, using a new FlowFISH method we have recently devised ^19^, we have unequivocally demonstrated that nucleated human RBCs have been infected *in vivo* (Extended data Fig. 2b), confirming previous observations by transmission electron microscopy of BM patients’ biopsies **^20^**. Fifth, the efficient parasite transmission from HIS-HEry mice with low parasite densities in PB to *Anopheles* mosquitoes recapitulates the observation that patients with very low parasitemia contribute to human-to-mosquito transmission and may constitute an important infectious reservoir **^21^.** Therefore, and in contrast with current mice model for human *Plasmodium* blood stage infection based on the adoptive transfer of human mature RBCs for P*. falciparum* **^22^** or reticulocytes for *P. vivax* **^23^**, the HIS-HEry model provides unique opportunities to address crucial questions associated with BM infection, such as parasite-induced dyserythropoiesis leading to anemia and more generally problems linked to hematopoietic disorders that may be associated with parasite-mediated inflammation **^6^**.

The observed increase in parasite load upon secondary transfer suggests that the selection of better adapted clinical isolates to the murine host environment could be achieved as it was previously shown for *P. falciparum* **^24^**. Multiple passages may result in higher parasitemia in our mouse model, which could provide the biomass for biochemical studies and even to establish transfection of *P. vivax* to explore for example drug resistance mechanisms in this pathogen.

The vast majority of infected HIS-HEry mice were able to transmit *P. vivax* to *Anopheles*, leading to production of salivary-gland sporozoites. This was also true for day 21 infected mice, showing that the production of infective transmission stages is maintained for at least 3 weeks. The experimental generation of *P. vivax* sporozoites currently requires fresh blood from infected *P. vivax* patients to be used to feed mosquitoes in *P. vivax* endemic regions, **^25^** while our HIS-HEry mice opens the possibility to produce *P. vivax* sporozoites in any malaria research center. Given that 2-100 sporozoites were enough to produce infect a human volunteer by mosquitoes fed with *P. vivax* infected patients’ blood ^26^, the HIS-HEry mice can provide sufficient amount of sporozoites for experimental infection of humans. The combination of primary human liver cell culture and *P. vivax* sporozoites will open avenues to study this neglected but important topic, including hypnozoite forms, which remains a major hurdle to disease eradication **^4^**.

In conclusion, the newly established humanized mouse model of long-lasting *P. vivax* infection should offer a unique opportunity for *in vivo* testing of novel antimalarial interventions. It will facilitate testing new vaccine strategies against asexual and sexual stages and provide a platform for *in vivo* screening of new blood schizonticides. Importantly it will represent a promising model to devise new transmission-blocking strategies both at the levels of vertebrate and invertebrate hosts.

**Extended Fig. 1 | HIS-HEry chimeras support efficient human erythropoiesis.** BM human erythrocytes from 16 to 24 weeks old HIS-HEry chimeras were analyzed by flowcytometry for the expression of: **a**-CD49d and Band3 surface markers shown as a representative dot plot; **b**, Intracellular mature human hemoglobin subunit beta (HBB) and fetal hemoglobin (HBF) shown as a representative dot plot. **c**, CD71, DARC (CD234), CD36 and CD44 shown as representative histograms (filled) compared to expression in control non-reconstituted mouse (open). d**-** Representative quantification of human CD235a^+^ RBCs in HIS-HEry PB without (Clod-) or under (Clod+) clodronate liposome treatment (3 times at 5 days intervals) (*n* = 4 mice).

**Extended Fig. 2 | HIS-HEry support sexual stages differentiation of *P vivax.* a**, ImageStream profile of both sexual and asexual stages in endogenous human RBCs (RhD+) from day7 infected HIS-HEry after FlowFISH staining. The staining are the following: RhD marker (FITC Chanel 02), Pvs25 (Quasar 570 – Cy3 like Chanel 03), DAPI (Chanel 07), Pv18S RNA (Quasay 670 – Cy5 like Chanel 11), CD235a (APC-vio 770 Chanel 12) and merge. **b**, Heat map numbers of Pvs25 RNA copy numbers per 10^5^ human CD235a^+^ BM cells from HIS-HEry chimeras at day1 and day7 post-infection determined by qRT-PCR. Individual mice (*n* = 4) are shown per day of analysis. Mean day1 = 0.85 ±1.30; mean day7 = 1.25 ± 1.76. *P*-value (one-sample *t*-test) = 0.737. Freedom degree = 3.

## Methods

### Generation of Rag2^tm1Fwa^ IL2rgt^m1Cgn^ B2m^tm1Unc^ H2-Ab1^tm1Doi^ Tg(HLA-DRA*0101,HLA-DRB1*0101)1Dma Tg(HLA-A2) Tg(SIRPA) Hc^0^ cKitW41^d/d^ (CH1-2hSaW41) host mice

CH1-2hSa mice generated in our laboratory ^8^ were crossed with the B6W41 mice (generously provided by Claudia Waskow, University of Dresden, Germany) to generate CH1-2hSaW41 mice. Mice were bred and maintained in dedicated facilities of the Institut Pasteur. Procedures involving mice were previously approved by local Animal Ethics Committees and registered with the French authorities.

### Isolation of CD34+ human cord blood HSPCs and transplantation into CH1-2hSaW41 host mice

Cord bloods were obtained from de-identified healthy donors (AP-HP, Hôpital Saint-Louis, Unité de Thérapie Cellulaire, CRB-Banque de Sang de Cordon, Paris, France – authorization number: AC-2016-2759). Mononuclear cells were enriched by Ficoll Hypaque followed by the positive selection of CD34^+^ cells by immunomagnetic bead selection using AutoMACS pro instrument (Myltenyi Biotec) and the purity (>95%) was checked by FACS. RhD expression were determined by Flowcytometry (see below). Alternatively, human cord blood CD34+ cells were purchased from AbcellBio company, France.

To generate HIS-HEry chimera, 1-4 days old CH1-2hSaW41 neonates were injected into the liver with 10-30 000 human cord blood CD34^+^ cells in 30μl of phosphate buffered saline (PBS) using a 30-gauge needle (BD) as previously described. For some experiments, mice were injected i.v. under anesthesia with 200μl clodronate liposomes and blood samples were collected one day after to assess the RhD status of human RBCs by flowcytometry (Cytometry section).

### Flow cytometry

For human erythrocytes cell surface analysis, cells were stained with anti-Ly5.2, CD45, Ter119, CD235a, CD49d, Band3, DAPI, CD71, CD234 (DARC), CD44 monoclonal or anti-RhD antibodies. Secondary anti-human IgG1 mAbs were used for revealing anti-RhD staining. For adult HbB and fetal HbF hemoglobin expression, cells were first stained with anti-Ly5.2, CD45, TER119 and CD235a then fixed for 30 min on 500uL of a 4% PFA solution (Thermo Scientific™) in PBS and 0,0075% glutaraldehyde (Electron Microscopy Sciences). For dead cell exclusion, cells were stained using the LIFE-DEAD™ fixable blue dead cell stain kit (Invitrogen) following the manufacturer’s instructions or 7AAD (Sigma). After permeabilization with Triton 1mM, they were co-stained with anti-HbB and HbF monoclonal antibodies and washed two times with PBS. Cells were processed on a BD LSRFortessa™ (BD Biosciences Becton Dickinson) and data were analyzed using FlowJo LLC software v10.6.1 (Becton Dickinson). All human RBCs were defined as Ly5.2^−^ TER119^−^ CD45^−^ CD235a^+^ cells.

### *P. vivax* isolates

Seven clinical *P. vivax* samples collected in northwestern Brazil in the context of an ongoing cohort study (ClinicalTrials.gov, NCT03689036) were leukocyte-depleted and cryopreserved in liquid nitrogen as described ^27^ (Supplemental Table 1). Study protocols have been approved by the Institutional Review Board of the Institute of Biomedical Sciences, University of São Paulo, and by the National Human Research Ethics Committee of the Ministry of Health of Brazil (CAAE: 64767416.6.0000.5467); all patients provided written informed consent. RhD expression was determined for each isolate as described in the flowcytometry section.

### HIS-HEry mice infection with *P. vivax*

*P. vivax* isolates were thawed using a stepwise NaCl method as previously reported ^28^, washed in PBS and tested for *Plasmodium* parasitemia before mice infection by thick smear and Giemsa staining. RBC pellets were resuspended into PBS at 2.5 10^6^ to 5.10^6^ parasitized RBCs per ml. One day before infection, 12 to 20 weeks old HIS-HEry chimeras were treated by i.v. injection of 200 ml liposome clodronate (Liposoma BV, The Netherlands), then each 5 days for all the duration of experiments. HIS-HEry mice were infected i.v. under anesthesia with 10^6^-5.10^6^ infected RBCs. BM and blood were collected on euthanized mice at d1, 7, 14 or 21 after infection. For BM, after dissection of the 2 femurs, tibias, and humerus, the bones were crushed, and cell suspensions were filtered through 40 μM filters before processing. Each experiment has been done on chimeras engrafted with CD34^+^ cells from the same CB donor.

### *P. vivax* RNA quantification and characterization by qRT-PCR

Dry ice frozen PB and 10^6^ BM cell pellet were resuspended in TRIzol LS reagent followed by RNA extraction as described by the manufacturer (Invitrogen, Carlsbad, CA). Twenty μl of cDNA was then synthesized using the single-tube procedure of the SuperScript™ II Reverse Transcriptase cDNA synthesis kit (Invitrogen Carlsbad, CA), according to the manufacturer’s instructions. Quantitative PCR was performed on 2 μl with the SsoAdvanced™ Universal SYBR^®^ Green Supermix (Bio-Rad) and the following primer pairs: CytB-Forward: 5’-TGGAGTGGATGGTGTTTTAGA 3’ and CytB-Reverse: 5’ TTGCACCCCAATAACTCATTT-3’; Pvs25-Forward: 5’-AACGAAGGGCTGGTGCACCTTT-3’ and Pvs25-Reverse: 5’-AGCAACCTGCACTTTGGATTTCCG-3’. In parallel, different dilutions of a plasmid solution containing 1 copy of *Plasmodium* CytB and Pvs25 (from 10^2^ to 10^5^ copies) were amplified in the same conditions to set up a standard curve. The data analysis was done using Bio-Rad CFX Maestro Version 4.1.2434.0124. Results were expressed numbers of CytB or Pvs25 RNA copies per 10^5^ human BM CD235a^+^ RBCs deduced after FACS analysis of the % of CD235a^+^ cells.

### *P. vivax* quantification and sexual characterization by FlowFISH

We recently described this method adapted from the commercial method Stellaris® ^19^. Both PB and BM were first stained with LIVE/DEAD™ Fixable Blue Dead Cell Stain Kit (Invitrogen) and conjugated anti-TER119, -CD235a, -CD71 mAbs and in some experiments non-conjugated anti-RhD mAbs followed by FITC conjugated anti-human IgG1. Stained cells were fixed for 30 min on 500uL of a 4% PFA solution (Thermo Scientific™) in PBS and 0,0075% glutaraldehyde (Electron Microscopy Sciences) at room temperature (RT), then washed two times with PBS. Fixed cells were hybridized following the manufacturer’s instructions (www.biosearchtech.com/stellarisprotocols) with a set of anti-*P. vivax* 18S rRNA quasar-670-conjugated and anti-*P. vivax* Pvs25 quasar-570-conjugated RNA Stellaris FISH Probes (Biosearch Technologies). RNA Probes were designed using the Stellaris® FISH Probe Designer (Biosearch Technologies) available online at www.biosearchtech.com/stellarisdesigner. Importantly, the Pv18S probes didn’t cross-reacted with mouse 18S RNA. For flow image procedure, hybridized RBCs were then stained with DAPI as described previously. Cells were processed on a BD LSRFortessa™ (BD Biosciences Becton Dickinson) and data were analyzed using FlowJo LLC software v10.6.1 (Becton Dickinson).

For sporozoites detection, salivary glands were collected from fed *Anopheles* mosquitoes, dilacerated and filtered on 35 μm mesh. After washing in PBS, recovered sporozoites cells were stained with anti-PvCSP mAbs 1:500 and fixed 10 min in 100uL of a 2% PFA solution (Thermo Scientific™) in PBS. Sporozoites were then stained with DAPI as described above. Sporozoites were acquired using an Image stream ISX MkII flow cytometer (Amnis Corp, EMD Millipore) with channel 01 for BrightField, channel 02 for FITC-conjugated anti-PvCSP and channel 07 for DAPI. Data were analyzed using the IDEAS software (version 6.2).

### Secondary transfers of BM cell from infected HIS-HEry into naïve HIS-HEry chimeras

BM human CD235a+ cells collected from infected HIS-HEry chimeras were first enriched by AutoMACS pro and FACS sorted. Cells were then either directly injected i.v. into clodronate liposome treated-secondary naïve HIS-HEry mice or frozen in 50% IMDM 50% freezing solution (SVF 20% DMSO). Frozen cells were thawed and injected (10 000 to 120 000 cells) into clodronate liposome treated-secondary naïve HIS-HEry mice as for fresh cells.

### Mosquito feeding

Female *Anopheles stephensi* were either commercially provided by Radboud University Medical Center (RUMC), Nijmegen, The Netherlands or reared in the Centre for Production and Infection of A*nopheles* (CEPIA) at Pasteur Institute using standard procedures and SDA500 strain. HIS-HEry infected mice were anesthetized i.p. with approximately 70 mg/kg of ketamine and 2 mg/kg xylazine before their transfer to the BSL-3 facility in CEPIA facility. Anesthesized mice were placed on a mesh screen covering the container of mosquitoes with 30-60 mosquitos per mouse. Mosquitoes were then allowed to feed for 10-15 min. Mice were then euthanized and BM and PB were collected for further analysis. Blood feeding was confirmed through visualization of blood engorged female mosquitoes. Unfed mosquitoes were removed, and the fed mosquitoes were maintained at the CEPIA facility, Institut Pasteur Paris under standard breeding conditions. Salivary glands from pooled mosquitos were dissected at 14 post-feeding and sporozoites were identified and quantified by FlowFISH followed by Amnis analysis.

## Acknowlegments

We are grateful to Maria José Menezes for expert help with parasite sample collection and processing. The anti-PvCSP MRA-184 mAbs from 2F2 hybridoma were obtained through BEI Resources, NIAID, NIH. We thank Sara El Hoss from the Université de Paris, UMR_S1134, BIGR, Inserm, Institut National de Transfusion Sanguine, F-75015 Paris France for Band3 / CD49d staining. We thank Prof. Yves Colin for the anti-human RhD mAbs F5 the Institut National de la Santé et de la Recherche Médicale (INSERM) U76, Institut National de la Transfusion Sanguine (INTS); and (D.D.) INSERM U409, Faculté de Médecine Xavier Bichat, Paris, France. We acknowledge the Center for Translational Science (CRT)-Cytometry and Biomarkers Unit of Technology and Service (CB UTechS) and CEPIA (Centre de production et d’infection des Anophèles) at Institut Pasteur for support in conducting this study. *Anopheles stephensi* mosquitoes have been obtained from Radboud University Medical Center (RUMC) Nijmegen. at Institut Pasteur for support in conducting this study. We thank Coralie Guerin and Anna Chipont from the Cytometry Core from Institut Curie for providing access to ImageStream.

## Author contribution

CLB, LSM, ST and SG conceived and designed experiments. CLB, ST, MSH, FN, AC and SG performed experiments. CLB and SG analyzed data. VCN and MUF collected and provided clinical samples. SG, AS, LMS and MUF wrote the manuscript. All authors reviewed the manuscript for intellectual content and approved the final version to be submitted for publication.

## Additional informations

Supplemental table describes *P. vivax* isolates, parasitemia and numbers of parasites injected into HIS-HEry chimeras.

## Main references

1 World Malaria Report, WHO, 2020, ISBN 978-92-4-001579-1, https://www.who.int/malaria..

2 Obaldia, N., 3rd et al. Bone Marrow Is a Major Parasite Reservoir in Plasmodium vivax Infection. MBio 9, doi:10.1128/mBio.00625-18 (2018).

3 Merrick, C. J. Hypnozoites in Plasmodium: Do Parasites Parallel Plants? Trends Parasitol 37, 273–282, doi:10.1016/j.pt.2020.11.001 (2021).

4 Price, R. N., Commons, R. J., Battle, K. E., Thriemer, K. & Mendis, K. Plasmodium vivax in the Era of the Shrinking P. falciparum Map. Trends Parasitol 36, 560–570, doi:10.1016/j.pt.2020.03.009 (2020).

5 Gruszczyk, J. et al. Transferrin receptor 1 is a reticulocyte-specific receptor for Plasmodium vivax. Science 359, 48–55, doi:10.1126/science.aan1078 (2018).

6 Silva-Filho, J. L. et al. Plasmodium vivax in Hematopoietic Niches: Hidden and Dangerous: (Trends in Parasitology 36, 447-458, 2020). Trends Parasitol 36, 648–649, doi:10.1016/j.pt.2020.05.006 (2020).

7 Gunalan, K., Rowley, E. H. & Miller, L. H. A Way Forward for Culturing Plasmodium vivax. Trends Parasitol 36, 512–519, doi:10.1016/j.pt.2020.04.002 (2020).

8 Serra-Hassoun, M. et al. Human hematopoietic reconstitution and HLA-restricted responses in nonpermissive alymphoid mice. J Immunol 193, 1504–1511, doi:10.4049/jimmunol.1400412 (2014).

9 Rahmig, S. et al. Improved Human Erythropoiesis and Platelet Formation in Humanized NSGW41 Mice. Stem Cell Reports 7, 591–601, doi:10.1016/j.stemcr.2016.08.005 (2016).

10 Cosgun, K. N. et al. Kit regulates HSC engraftment across the human-mouse species barrier. Cell Stem Cell 15, 227–238, doi:10.1016/j.stem.2014.06.001 (2014).

11 McIntosh, B. E. et al. Nonirradiated NOD,B6.SCID Il2rgamma-/-Kit(W41/W41) (NBSGW) mice support multilineage engraftment of human hematopoietic cells. Stem Cell Reports 4, 171–180, doi:10.1016/j.stemcr.2014.12.005 (2015).

12 Wangen, J. R., Eidenschink Brodersen, L., Stolk, T. T., Wells, D. A. & Loken, M. R. Assessment of normal erythropoiesis by flow cytometry: important considerations for specimen preparation. Int J Lab Hematol 36, 184–196, doi:10.1111/ijlh.12151 (2014).

13 Horuk, R. et al. A receptor for the malarial parasite Plasmodium vivax: the erythrocyte chemokine receptor. Science 261, 1182–1184, doi:10.1126/science.7689250 (1993).

14 Yurino, A. et al. Enhanced Reconstitution of Human Erythropoiesis and Thrombopoiesis in an Immunodeficient Mouse Model with Kit(Wv) Mutations. Stem Cell Reports 7, 425–438, doi:10.1016/j.stemcr.2016.07.002 (2016).

15 Luiza-Batista, C. et al. Flowcytometric and ImageStream RNA-FISH gene expression, quantification and phenotypic characterization of blood and liver stages from human malaria species. The Journal of Infectious Diseases in press (2021).

16 Sa, J. M., Cannon, M. V., Caleon, R. L., Wellems, T. E. & Serre, D. Single-cell transcription analysis of Plasmodium vivax blood-stage parasites identifies stage- and species-specific profiles of expression. PLoS Biol 18, e3000711, doi:10.1371/journal.pbio.3000711 (2020).

17 Baro, B. et al. Plasmodium vivax gametocytes in the bone marrow of an acute malaria patient and changes in the erythroid miRNA profile. PLoS Negl Trop Dis 11, e0005365, doi:10.1371/journal.pntd.0005365 (2017).

18 Adapa, S. R. et al. Plasmodium vivax readiness to transmit: implication for malaria eradication. BMC Syst Biol 13, 5, doi:10.1186/s12918-018-0669-4 (2019).

19 Luiza-Batista, C. et al. Flowcytometric and ImageStream RNA-FISH gene expression, quantification and phenotypic characterization of blood and liver stages from human malaria species. J Infect Dis, doi:10.1093/infdis/jiab431 (2021).

20 Ru, Y. X. et al. Invasion of erythroblasts by Pasmodium vivax: A new mechanism contributing to malarial anemia. Ultrastruct Pathol 33, 236–242, doi:10.3109/01913120903251643 (2009).

21 Alves, F. P. et al. Asymptomatic carriers of Plasmodium spp. as infection source for malaria vector mosquitoes in the Brazilian Amazon. J Med Entomol 42, 777–779, doi:10.1093/jmedent/42.5.777 (2005).

22 Tyagi, R. K. et al. Humanized Mice Are Instrumental to the Study of Plasmodium falciparum Infection. Front Immunol 9, 2550, doi:10.3389/fimmu.2018.02550 (2018).

23 Schafer, C. et al. A Humanized Mouse Model for Plasmodium vivax to Test Interventions that Block Liver Stage to Blood Stage Transition and Blood Stage Infection. iScience 23, 101381, doi:10.1016/j.isci.2020.101381 (2020).

24 Jimenez-Diaz, M. B. et al. Improved murine model of malaria using Plasmodium falciparum competent strains and non-myelodepleted NOD-scid IL2Rgammanull mice engrafted with human erythrocytes. Antimicrob Agents Chemother 53, 4533–4536, doi:10.1128/AAC.00519-09 (2009).

25 Collins, K. A. et al. A Plasmodium vivax experimental human infection model for evaluating efficacy of interventions. J Clin Invest 130, 2920–2927, doi:10.1172/JCI134923 (2020).

26 Herrera, S. et al. Consistent safety and infectivity in sporozoite challenge model of Plasmodium vivax in malaria-naive human volunteers. Am J Trop Med Hyg 84, 4–11, doi:10.4269/ajtmh.2011.09-0498 (2011).

## Methods references

27 de Oliveira, T. C. et al. Genome-wide diversity and differentiation in New World populations of the human malaria parasite Plasmodium vivax. PLoS Negl Trop Dis 11, e0005824, doi:10.1371/journal.pntd.0005824 (2017).

28 Saunders, G. M., Talmage, D. W. & Scott, V. The use of Plasmodium vivax preserved by freezing in inducing malaria. J Lab Clin Med 33, 1579–1587 (1948).

