## Supplementary figures and images for "Humanized mice for sustained *Plasmodium vivax* blood-stage infection and transmission"

### Supplemental Figure 1

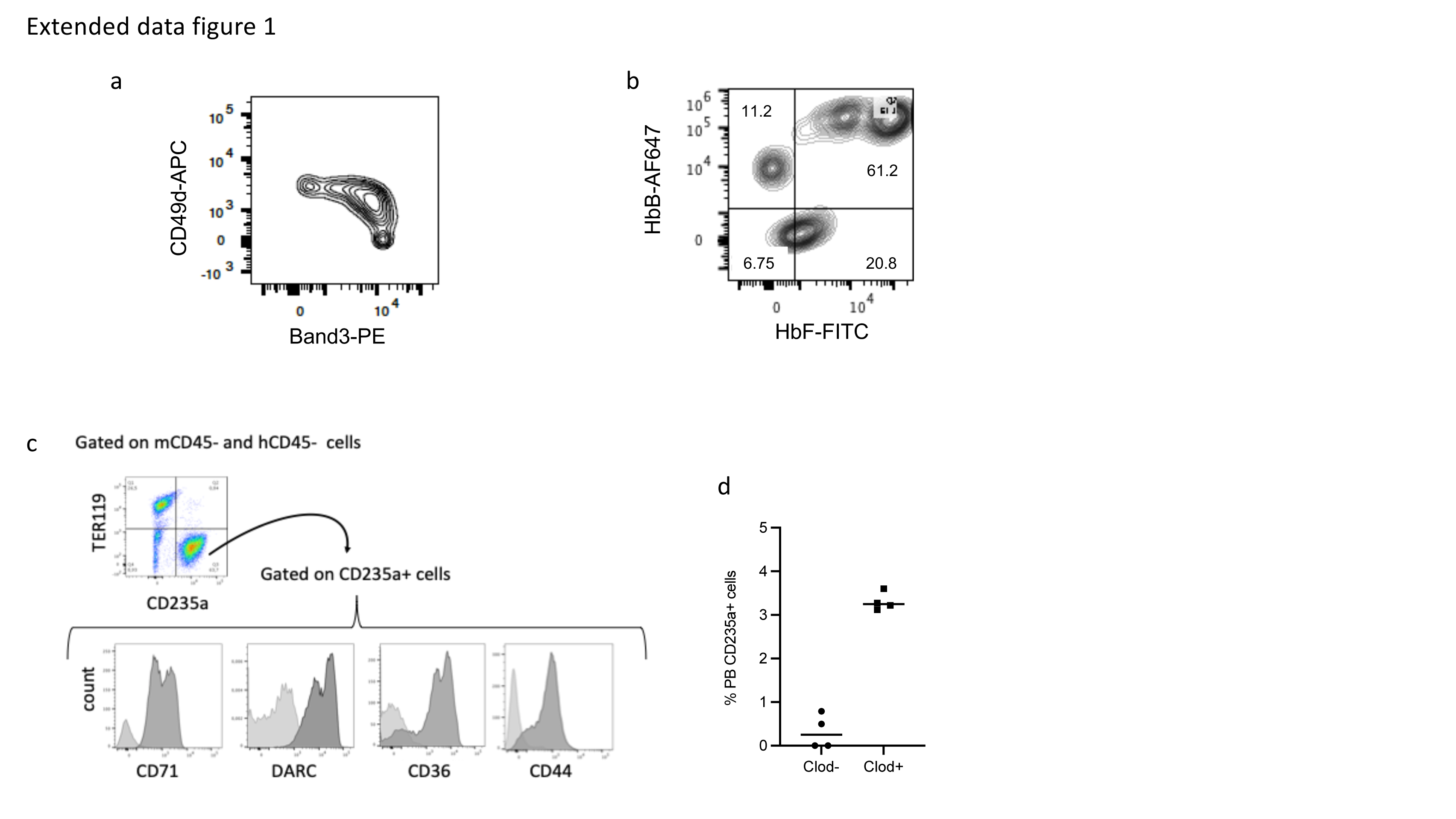

### Supplemental Figure 2

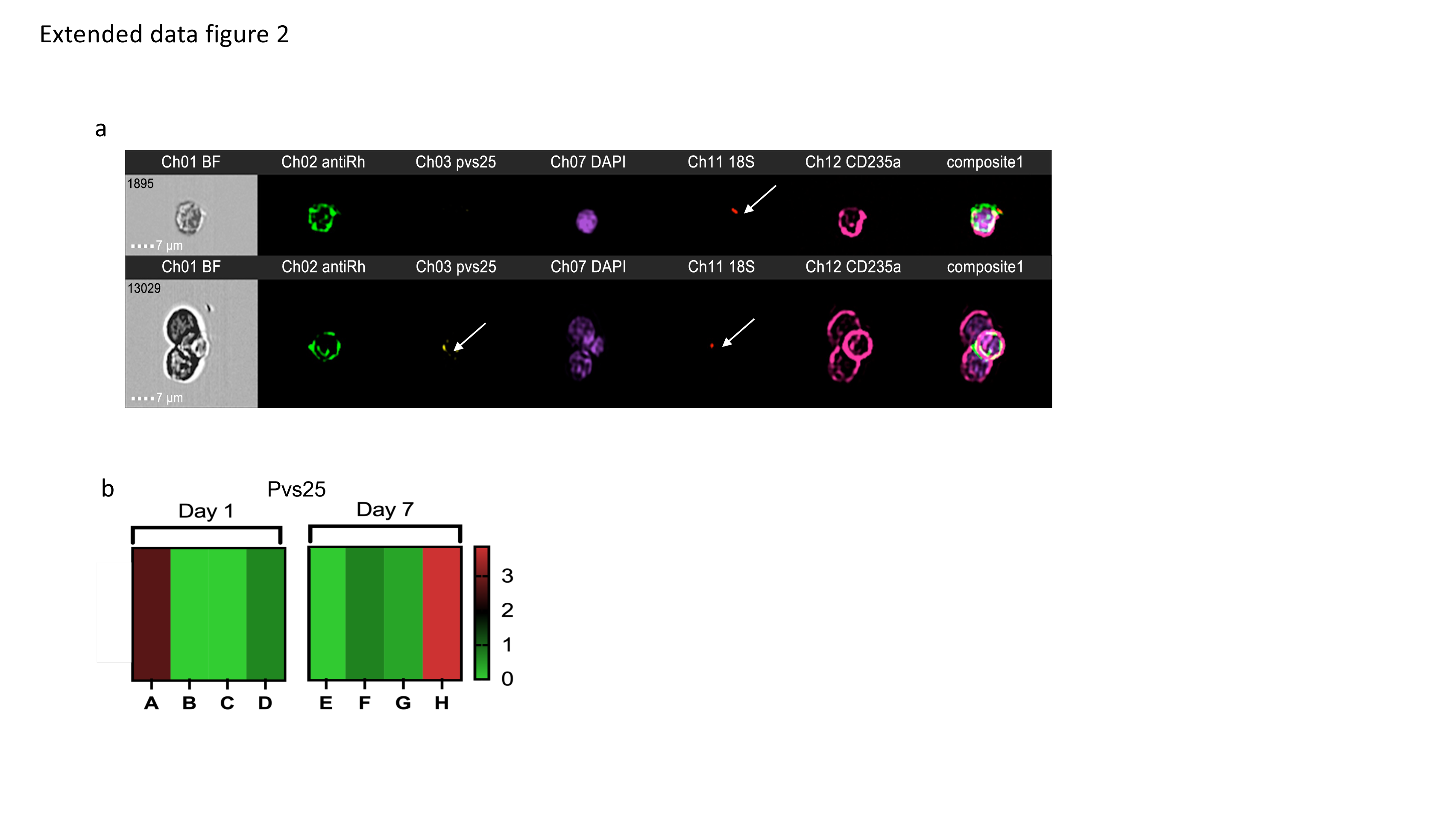
