## Supplemental table for "Humanized mice for sustained *Plasmodium vivax* blood-stage infection and transmission"

| # *Pv* isolate | parasitemia (%) | # parasites injected  per mouse |
| --- | --- | --- |
| Pv1 | 0.045 | 9 x 10^5^ |
| Pv2 | 0.031 | 6.2 x 10^5^ |
| Pv3 | 0.023 | 4.6 x 10^5^ |
| Pv4 | 0.015 | 3 X 10^5^ |
| Pv5.1  Pv5.2 Pv5  Pv5.3 | 0.044  0.030 0.035  0.031 | 2.9 x 10^5^  1 x 10^5^ 4.9 X 10^5^  1 x 10^5^ |

***P. vivax* (Pv) isolates from Acre (Brazil) used to experimentally infect HIS-HEry chimeric mice**. Mice were inoculated with either a single isolate (Pv1, Pv2, Pv3, and Pv4) or a mixture of 3 isolates (Pv5), except if stated otherwise in figure legends. Percent parasitemia (measured by microscopy of Giemsa-stained blood smears) and the number of infected blood cells per inoculum are shown; parasitemia for Pv5 is an average of those in individual isolates. All isolates tested negative for *P. falciparum* by qPCR.
